# Relationship Between Prediction Accuracy and Feature Importance Reliability: an Empirical and Theoretical Study

**DOI:** 10.1101/2022.08.08.503167

**Authors:** Jianzhong Chen, Leon Qi Rong Ooi, Trevor Wei Kiat Tan, Shaoshi Zhang, Jingwei Li, Christopher L. Asplund, Simon B Eickhoff, Danilo Bzdok, Avram J Holmes, B.T. Thomas Yeo

## Abstract

There is significant interest in using neuroimaging data to predict behavior. The predictive models are often interpreted by the computation of feature importance, which quantifies the predictive relevance of an imaging feature. Tian and Zalesky (2021) suggest that feature importance estimates exhibit low split-half reliability, as well as a trade-off between prediction accuracy and feature importance reliability across parcellation resolutions. However, it is unclear whether the trade-off between prediction accuracy and feature importance reliability is universal. Here, we demonstrate that, with a sufficient sample size, feature importance (operationalized as Haufe-transformed weights) can achieve fair to excellent split-half reliability. With a sample size of 2600 participants, Haufe-transformed weights achieve average intra-class correlation coefficients of 0.75, 0.57 and 0.53 for cognitive, personality and mental health measures respectively. Haufe-transformed weights are much more reliable than original regression weights and univariate FC-behavior correlations. Original regression weights are not reliable even with 2600 participants. Intriguingly, feature importance reliability is strongly positively correlated with prediction accuracy across phenotypes. Within a particular behavioral domain, there is no clear relationship between prediction performance and feature importance reliability across regression models. Furthermore, we show mathematically that feature importance reliability is necessary, but not sufficient, for low feature importance error. In the case of linear models, lower feature importance error is mathematically related to lower prediction error. Therefore, higher feature importance reliability might yield lower feature importance error and higher prediction accuracy. Finally, we discuss how our theoretical results relate with the reliability of imaging features and behavioral measures. Overall, the current study provides empirical and theoretical insights into the relationship between prediction accuracy and feature importance reliability.

## 1. Introduction

Neuroimaging provides a non-invasive means to study human brain structure and function. *In vivo* imaging features have been linked to many clinically relevant phenotypes when contrasting populations of patients and healthy controls (Greicius *et al*. 2004, Kennedy *et al*. 2006). However, these group-level studies ignore inter-individual differences within and across patient populations (Zhang *et al*. 2016, Xia *et al*. 2018, Zabihi *et al*. 2019, Tang *et al*. 2020, Wolfers *et al*. 2020). As a result, there is an increasing interest in the field to shift from group differences to accurate individual-level predictions (Dosenbach *et al*. 2010, Finn *et al*. 2015, Hsu *et al*. 2018, Nostro *et al*. 2018, Kong *et al*. 2019).

One goal of neuroimaging-based behavioral prediction is clinical usage to forecast practically useful clinical endpoints (Gabrieli *et al*. 2015). This ambition requires users to have trust in the predictive models, which often rests on a given models’ interpretability (Bussone *et al*. 2015, Price 2018, Anderson and Anderson 2019, Diprose *et al*. 2020, Hedderich and Eickhoff 2020). Indeed, the recently enacted European Union Global Data Protection Regulation (GDPR) states that patients have a right to “meaningful information about the logic involved” when automated decision-making systems are used (Vasey *et al*. 2022a, 2022b). Furthermore, in many studies, the derived predictive models are often interpreted to gain insights into the predictive principles and inter-individual differences that underpin observed brain-behavior relationships (Finn *et al*. 2015, Greene *et al*. 2018, Chen *et al*. 2022). Therefore, while many studies in the neuroimaging literature have focused on prediction accuracy (Dadi *et al*. 2019, He *et al*. 2020, Pervaiz *et al*. 2020, Schulz *et al*. 2020, Abrol *et al*. 2021), enhancing model interpretability remains an important issue.

One approach to interpret predictive models is the computation of feature-level importance, which quantifies the relevance of an imaging feature in the predictive model. In the case of linear models, most previous studies have interpreted the regression weights (Jiang *et al*. 2020, Sripada *et al*. 2020, Cropley *et al*. 2021, Xiao *et al*. 2021) of predictive models. However, the covariance structure among predictive features can lead to incorrect interpretations (Haufe *et al*. 2014). Instead, Haufe and colleagues demonstrated that it is necessary to perform an inversion of the linear models to yield the correct interpretation (Haufe *et al*. 2014). We refer to this inversion as the Haufe transform. Further explanation of the Haufe transform can be found in Section 2.7 and in the original study (Haufe *et al*. 2014).

A recent study suggested that in the context of behavioral predictions from functional connectivity (FC), the reliability of feature-level importance (original regression weights and Haufe-transformed weights) across independent samples was poor (Tian and Zalesky 2021). Because the study utilized a maximum sample size of 400 and predicted only a small selection of cognitive measures and sex, it remains unclear whether the results generalize to other sample sizes and behavioral domains. Tian and Zalesky also found that higher resolution parcellations led to better prediction accuracy but lower feature importance reliability. However, it is unclear whether the trade-off between prediction accuracy and feature importance reliability is universal. A universal trade-off would be counter-intuitive given that both feature importance reliability and prediction accuracy should reflect the reliability of brain-behavior relationship across independent datasets. More specifically, if the brain-behavior relationships in two independent data samples are highly similar, then we would expect that a model trained on one dataset to generalize well to the other dataset (i.e., high prediction accuracy). We would also expect the models trained on both datasets to be highly similar, leading to high feature importance reliability. Therefore, we hypothesize that there is not a universal trade-off between prediction accuracy and feature importance reliability.

In the present study, we used the Adolescent Brain Cognitive Development (ABCD) study to investigate the relationship between prediction accuracy and feature importance reliability. Resting-state functional connectivity was used to predict a wide range of 36 behavioral measures across cognition, personality (related to impulsivity), and mental health. We considered four commonly used prediction models: kernel ridge regression (KRR), linear ridge regression (LRR), least absolute shrinkage and selection operator (LASSO), and Random Forest (RF) models. Consistent with Tian and Zalesky (2021), we found that Haufe-transformed weights were more reliable than regression weights and univariate FC-behavior correlations. However, for sufficiently large sample sizes, we found fair to excellent split-half reliability for the Haufe-transformed weights. On the other hand, the original regression weights were unreliable even with thousands of participants. Intriguingly, feature importance reliability was strongly correlated with prediction accuracy across behavioral measures. Within a particular behavioral domain, there was no clear relationship between prediction performance and feature importance reliability across regression algorithms. We show mathematically that split-half feature importance reliability is necessary, but not sufficient, for low feature importance error. In the case of linear models, prediction error closely reflects feature importance error. Overall, the current study provides empirical and theoretical insights into the relationship between prediction accuracy and feature importance reliability.

## 2. Methods

### 2.1 Dataset

The Adolescent Brain Cognitive Development (ABCD) dataset (2.0.1 release) was used for its large sample size, as well as its rich imaging and behavioral measures. The Institutional Review Board (IRB) at the University of California, San Diego approved all aspects of the ABCD study (Auchter *et al*. 2018). Parents or guardians provided written consent while the child provided written assent (Clark *et al*. 2018).

After quality control and excluding siblings, the final sample consisted of 5260 unrelated participants. Consistent with our previous studies (Chen *et al*. 2022, Ooi *et al*. 2022), each participant had a 419 × 419 FC matrix as the imaging features, which were used to predict 36 behavioral measures across the behavioral domains of cognition, personality, and mental health.

### 2.2 Image preprocessing

Images were acquired across 21 sites in the United States with harmonized imaging protocols for GE, Philips, and Siemens scanners (Casey *et al*. 2018). We used structural T1 and resting-fMRI. For each participant, there were four resting-fMRI runs. Each resting-fMRI run was 300 seconds long. Preprocessing followed our previously published study (Chen *et al*. 2022). For completeness, the key preprocessing steps are summarized here.

Minimally preprocessed T1 data were used (Hagler *et al*. 2019). The structural data were further processed using FreeSurfer 5.3.0 (Dale *et al*. 1999, Fischl, Sereno, and Dale 1999, Fischl, Sereno, Tootell, *et al*. 1999, Fischl *et al*. 2001, Ségonne *et al*. 2004, 2007), which generated accurate cortical surface meshes for each individual. Individuals’ cortical surface meshes were registered to a common spherical coordinate system (Fischl, Sereno, and Dale 1999, Fischl, Sereno, Tootell, *et al*. 1999). Individuals who did not pass recon-all quality control (Hagler *et al*. 2019) were removed.

Minimally preprocessed fMRI data (Hagler *et al*. 2019) were further processed with the following steps: (1) removal of initial frames, with the number of frames removed depending on the type of scanner (Hagler *et al*. 2019); and (2) alignment with the T1 images using boundary-based registration (Greve and Fischl 2009) with FsFast (http://surfer.nmr.mgh.harvard.edu/fswiki/FsFast). Functional runs with boundary-based registration (BBR) costs greater than 0.6 were excluded. Framewise displacement (FD) (Jenkinson *et al*. 2002) and voxel-wise differentiated signal variance (DVARS) (Power *et al*. 2012) were computed using fsl_motion_outliers. Respiratory pseudomotion was filtered out using a bandstop filter (0.31-0.43 Hz) before computing FD (Power *et al*. 2019, Fair *et al*. 2020, Gratton *et al*. 2020). Volumes with FD > 0.3 mm or DVARS > 50, along with one volume before and two volumes after, were marked as outliers and subsequently censored. Uncensored segments of data containing fewer than five contiguous volumes were also censored (Gordon *et al*. 2016, Kong *et al*. 2019). Functional runs with over half of their volumes censored and/or max FD > 5mm were removed. Individuals who did not have at least 4 minutes of data were also excluded from further analysis.

The following nuisance covariates were regressed out of the fMRI time series: global signal, six motion correction parameters, averaged ventricular signal, averaged white matter signal, and their temporal derivatives (18 regressors in total). Regression coefficients were estimated from the non-censored volumes. We chose to regress the global signal because we were interested in behavioral prediction, and global signal regression has been shown to improve behavioral prediction performance (Greene *et al*. 2018, Li *et al*. 2019). The brain scans were interpolated across censored frames using least squares spectral estimation (Power *et al*. 2014), band-pass filtered (0.009 Hz ≤ f ≤ 0.08 Hz), projected onto FreeSurfer fsaverage6 surface space and smoothed using a 6 mm full-width half maximum kernel.

### 2.3 Functional connectivity

We used a whole-brain parcellation comprising 400 cortical regions of interest (ROIs) (Schaefer *et al*. 2018) and 19 subcortical ROIs (Fischl *et al*. 2002). For each participant and each fMRI run, functional connectivity (FC) was computed as the Pearson’s correlations between the average time series of each pair of ROIs. FC matrices were then averaged across runs, yielding a 419□×□419 FC matrix for each participant. Correlation values were converted to z-scores using Fisher’s r-to-z transformation prior to averaging and converted back to correlation values after averaging. Censored frames were ignored when computing FC.

### 2.4 Behavioral data

Following our previous study (Chen *et al*. 2022), we considered 16 cognitive, 11 mental health, and 9 impulsivity-related personality measures. The cognitive measures were vocabulary, attention, working memory, executive function, processing speed, episodic memory, reading, fluid cognition, crystallized cognition, overall cognition, short delay recall, long delay recall, fluid intelligence, visuospatial accuracy, visuospatial reaction time, and visuospatial efficiency. The mental health measures were anxious depressed, withdrawn depressed, somatic complaints, social problems, thought problems, attention problems, rule-breaking behavior, aggressive behavior, total psychosis symptoms, psychosis severity, and mania. The impulsivity-related personality measures were negative urgency, lack of planning, sensation seeking, positive urgency, lack of perseverance, behavioral inhibition, reward responsiveness, drive, and fun seeking.

Participants who did not have all behavioral measures were excluded from further analysis. As recommended by the ABCD consortium, individuals from Philips scanners were also excluded due to incorrect preprocessing. Finally, by excluding siblings, the main analysis utilized data from 5260 unrelated children.

### 2.5 Split-half cross-validation

ABCD is a multi-site dataset. To reduce sample size variability across sites, smaller sites were combined to create 10 “site-clusters”, each containing at least 300 individuals (Table S1). Thus, participants within a site were in the same site-cluster.

A split-half cross-validation procedure was utilized to evaluate the prediction performance and the split-half reliability of feature importance. For each split, 5 site-clusters were selected as the training set and the remaining 5 were selected as the test set. Prediction models were trained on the training set to predict the behavioral measures from the FC matrices. The prediction models were then evaluated on the test set.

Here, we considered kernel ridge regression (KRR), linear ridge regression (LRR), and least absolute shrinkage and selection operator (LASSO) models for prediction. Hyperparameters were tuned using cross-validation within the training set (Chen et al., 2022). We also explored the use of random forests (RF) for prediction (Breiman 2001). Because of the large number of FC features in the current study, the RF is much slower than KRR: a single RF model required 2 hours of training compared with 10 seconds for KRR. Therefore, tuning the hyperparameters of the RFs would be computational infeasible. Consequently, the hyperparameters of the RF models were fixed with the number of trees set to 100 and the depth of each tree set to be 4.

Prediction accuracy was defined as the Pearson’s correlation between the predicted and observed behavior of test participants. Feature importance of the regression models was computed in the training set (see Section 2.7). After the prediction model was trained and evaluated, the training and test sets were swapped. The model training and evaluation procedure were then repeated. Thus, for a given regression approach and interpretation method, each data split yielded two prediction accuracies and two sets of feature importance.

For each data split, the two accuracy numbers were averaged yielding an overall prediction accuracy for the split. On the other hand, the two sets of feature importance (*f*_l_ and *f*_2_) were used to compute split-half reliability (Tian and Zalesky 2021), which we refer to as split-half intra-class correlation coefficient (ICC). Note that *f*_l_ and *f*_2_ are vectors of length *K*, where *K* is the total number of features. The *k*-th element of *f*_l_ (or *f*_2_) is the feature importance of the *k*-th feature in the first (or second) set of feature importance.

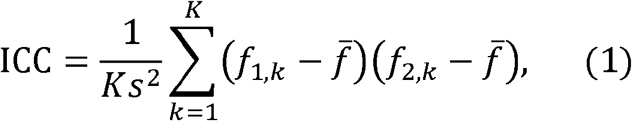

where *f*_l,*k*_ and *f*_2,*k*_ are the *k*-th element of *f*_l_ and *f*_2_ respectively. 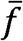 is the pooled mean computed from both *f*_1_ and *f*_2_, given by 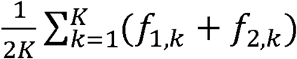. *s*^2^ is the pooled variance computed from both *f*_1_ and *f*_2_, given by 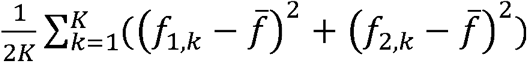.

To ensure stability, the data split was repeated 126 (the number of unique ways to split ten site-clusters into two halves, which is 10 choose 5 divided by 2) times.

### 2.6 Reliability across different sample sizes

The procedure in the previous section utilized the full sample size. To evaluate feature importance reliability across different sample sizes, the previous procedure (Section 2.5) was repeated, but the participants were subsampled for each split-half cross-validation to achieve a desired sample size *N*. More specifically, we considered sample sizes of 200, 400, 1000, and 1500. For each sample size *N*, we first split the 10 site-clusters into two halves, each containing 5 site-clusters (Section 2.5). *N*/10 samples were then randomly sampled from each site-cluster. The procedure was repeated 126 (the number of unique ways to split ten site-clusters into two halves, which is 10 choose 5 divided by 2) times.

### 2.7 Original and Haufe-transformed weights

We used KRR, LRR, LASSO and random forests (RF) to predict 36 behavioral measures from FC features. In particular, the lower triangular entries of the FC matrix were used as input for the regression models. LRR, LASSO and RF are commonly used in the literature. We have previously demonstrated that KRR is a powerful approach for resting-FC behavioral prediction (He *et al*. 2020).

Since KRR is less commonly used in the literature, we will provide a high-level explanation here. Briefly, let *y*_*i*_ and *FC*_*i*_ be the behavioral measure and FC of training individual *i*. Let *y*_*t*_ and *FC*_*t*_ be the behavioral measure and FC of a test individual. Then, kernel regression would predict the test individual’s behavior as the weighted average of the training individuals’ behavior, i.e. *y*_*t*_ ≈ Σ_*i*∈*tranining set*_ *similarity* (*FC*_*i*_,*FC*_*t*_)*y*_*i*_, where *similarity* (*FC*_*i*_,*FC*_*t*_) was defined as the Pearson’s correlation between *FC*_*i*_ and *FC*_*t*_. Thus, kernel regression assumed that individuals with more similar FC exhibit more similar behavior. To reduce overfitting, an l_2_-regularization term was included, which was tuned in the training set (Kong *et al*. 2019, Li *et al*. 2019, He *et al*. 2020).

To interpret the trained models, we considered both the regression weights and Haufe-transformed weights. Since LRR and LASSO are linear models, the regression weights were straightforward to obtain. In the case of KRR, the kernel regression model was converted to an equivalent linear regression model, yielding one regression weight for each feature (Liu *et al*. 2007, Chen *et al*. 2022). We note that this conversion was possible because we used the correlation kernel, which is linear when the input features are pre-normalized. In the case of RF models, feature importance was extracted through calculating the out-of-bag error using a conditional permutation procedure, which reduced selection bias of correlated variables (Strobl *et al*. 2008). We refer to this approach as conditional variable importance.

Each prediction model was also inverted using the Haufe transform (Haufe *et al*. 2014). To motivate the Haufe transform (Haufe et al., 2014; Chen et al., 2022), suppose we seek to predict target variable *y* (e.g., fluid intelligence) from the FC of two edges (*FC*1 and *FC*2). In this example, let us assume that *FC*1 = *y* − *motoin*, and *FC*2 = *motoin* Then a prediction model with 100% accuracy would be 1 × *FC*1 + 1 × *FC*2. The regression weights of this model are both one for *FC*1 and *FC*2. Based on the weights of the regression model, we would conclude that both *FC*1 and *FC*2 are strongly related to the target variable *y*. Haufe transform resolves this issue by computing the covariance between the predicted target variable and each FC feature in the training set. In this toy example, FC2 will be assigned a weight of zero by the Haufe transform, consistent with the intuition that FC2 is not related to the target variable even though it is helpful for predicting the target variable.

Further informative examples can be found in Haufe et al. (2014). More generally, Haufe and colleagues demonstrated that for a linear predictive model, the appropriate transformation can be obtained by computing the covariance of each feature and the predicted target variable in the training set. When applied to nonlinear models, the Haufe transform recovers the “best” linear interpretation of the nonlinear models in the least square sense (Haufe et al., 2014). Therefore, our application of the Haufe transform to the random forests will only yield a partial (linear) interpretation of the random forests.

### 2.8 Mass univariate associations

Besides predictive models, we also examined the split-half reliability of mass univariate associations between FC and behavioral measures, which is sometimes referred to as brain-wide association analysis (Marek *et al*. 2022). We note that mass univariate associations are often used for feature selection in neuroimaging predictive models (Finn *et al*. 2015). The selected features are then used to interpret the model (Finn *et al*. 2015, Shen *et al*. 2017). Therefore, mass univariate associations are a good proxy for such approaches. Here, univariate association is defined as the correlation between each FC feature and each behavioral measure. To study the split-half reliability of univariate associations, we performed the same split-half procedure (Sections 2.5 and 2.6). However, instead of training a predictive model in the training set, we correlated the FC features and the behavioral measures of the training participants to obtain one t-statistic for each feature and each behavioral measure. This procedure was repeated for the test participants. Split-half reliability was defined as the split-half ICC of the t-statistic values between the two halves of the dataset (i.e., training and test sets).

### 2.9 Data and code availability

The ABCD data are publicly available via the NIMH Data Archive (NDA). Processed data from this study have been uploaded to the NDA. Researchers with access to the ABCD data will be able to download the data: <u>LINK_TO_BE_UPDATED</u>. Analysis code specific to this study was can be found on GitHub: <u>LINK_TO_BE_UPDATED</u>. Co-authors NW and SZ reviewed the code before merging it into the GitHub repository to reduce the chance of coding errors.

## 3. Results

### 3.1 Haufe-transformed weights exhibit fair to excellent split-half reliability with large sample sizes

We computed resting-state functional connectivity (RSFC) among 400 cortical (Schaefer *et al*. 2018) and 19 subcortical (Fischl *et al*. 2002) regions for 5260 participants from the ABCD dataset (Casey *et al*. 2018). The lower triangular entries of the 419 × 419 RSFC matrix were then vectorized to predict 36 behavioral scores that span across 3 domains: cognition, personality, and mental health.

Feature importance of KRR predictive models was interpreted using two approaches: regression weights and Haufe-transformed weights. For comparison, t-statistics from mass univariate associations were also computed. We used a split-half procedure to compute the split-half reliability of feature importance. For each split, we fitted the KRR model on each half and obtained the feature importance. The split-half reliability was defined as the split-half ICC of the feature importance values between the two halves.

Figure 1 shows the split-half reliability of the two interpretation methods and mass univariate associations across 126 splits for different sample sizes and behavioral domains. Consistent with previous studies, split-half reliability of feature importance increases with larger sample sizes across all behavioral domains and interpretation methods (Tian and Zalesky 2021, Marek *et al*. 2022). The Haufe-transformed weights were consistently more reliable than univariate associations (t-statistics), which were in turn more reliable than the regression weights. Haufe-transformed weights at a sample size of 200 were more reliable than the original regression weights at a sample size of 2630.

**Figure 1.**
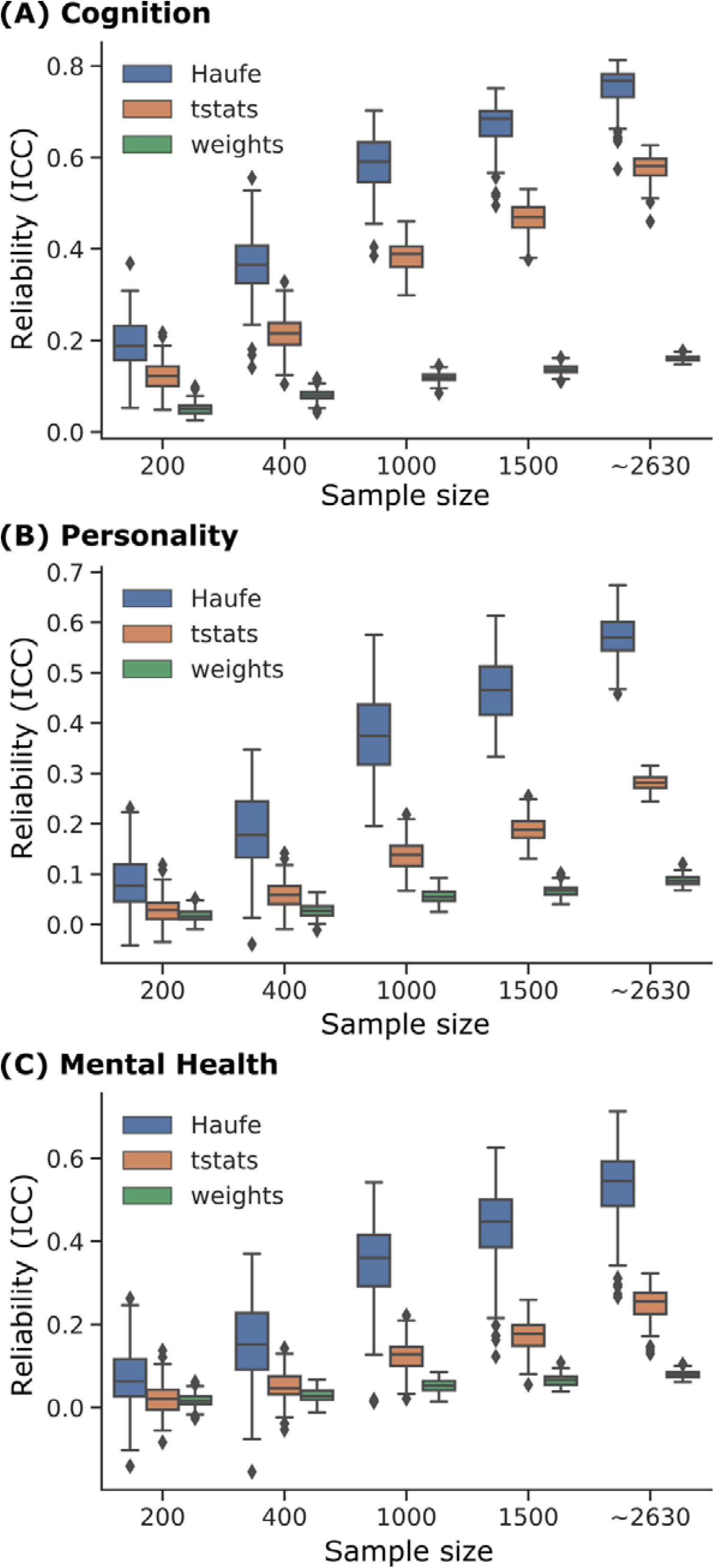
Split-half reliability of feature importance of kernel ridge regression (KRR) models across different sample sizes, interpretation methods, and behavioral domains: (A) cognition, (B) personality, and (C) mental health. Split-half reliability was computed as split-half interclass correlation coefficients (ICC) of feature importance obtained from two non-overlapping split-halves of the ABCD participants. After splitting, participants were randomly subsampled to show the effect of sample size on feature importance reliability. Full data without subsampling was reported as a sample size of ∼2630. “∼” was used because the two halves have similar (but not exactly the same) sample sizes that summed to 5260 (total number of participants). Split-half ICC values were reported for Haufe-transformed model weights (Haufe), mass univariate associations (tstats), and original regression weights (weights). Boxplots show the distribution of average split-half ICC within each behavioral domain across 126 split-half pairs. For each boxplot, the box extends from the lower to upper quartile values of the data, with a line at the median. The whiskers extend from the box to show the data range (excluding outliers). Outliers are defined as data points beyond 1.5 times the interquartile range and shown as flier points past the whiskers. Overall, across different sample sizes and behavioral domains, Haufe-transformed weights were more reliable than mass univariate associations (tstats), which were in turn more reliable than regression weights. Similar conclusions were obtained with linear ridge regression (Figure 2), LASSO (Figure S1) and random forests (Figure S2).

At the largest sample size of 2630, an average split-half ICC of 0.75 was achieved for Haufe-transformed weights of models predicting cognitive measures, which is considered “excellent” split-half reliability (Cicchetti 1994). On the other hand, an average split-half ICC of 0.57 and 0.53 were achieved for personality and mental health at the full sample size, which are considered “fair” split-half reliabilities (Cicchetti 1994). Under the same sample size and interpretation method, the split-half reliability of feature importance for mental health and personality was consistently lower than that of cognition.

Similar conclusions were obtained with linear ridge regression (Figure 2), LASSO (Figure S1) and RF (Figure S2). In the case of RF models, conditional variable importance was computed (instead of weights). Note that univariate associations (tstats) were computed independent of regression models and are therefore the same across Figures 1, 2, S1 and S2. Overall, we found that Haufe-transformed weights achieved fair to excellent split-half reliability with sufficiently large samples.

**Figure 2.**
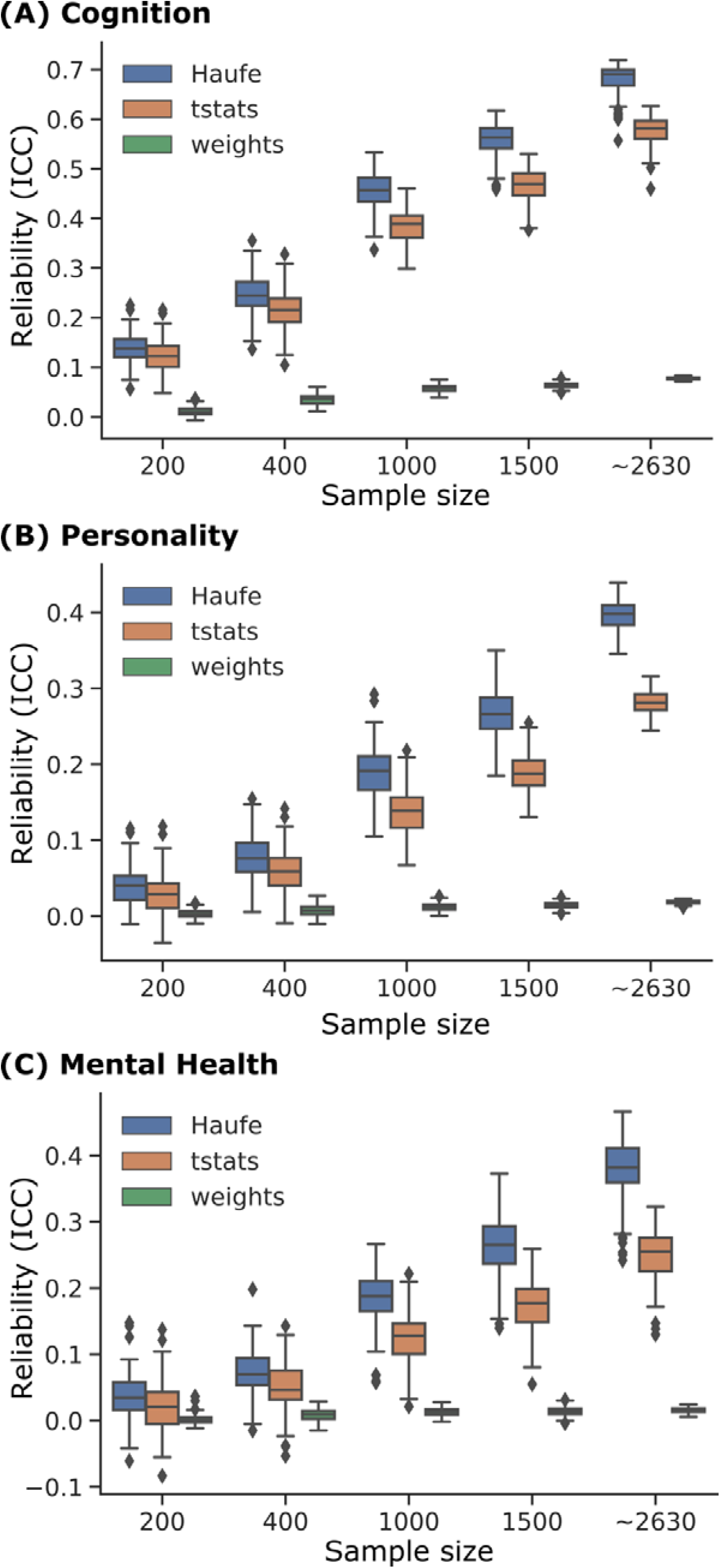
Split-half reliability of feature importance of linear ridge regression (LRR) models across different sample sizes, interpretation methods, and behavioral domains: (A) cognition, (B) personality, and (C) mental health. Same as Figure 1, except using LRR as the prediction model. Split-half reliability was computed as split-half interclass correlation coefficients (ICC) of feature importance obtained from two non-overlapping split-halves of the dataset. After splitting, data were randomly subsampled to show the effect of sample size on feature importance reliability. Full data without subsampling was reported as a sample size of ∼2630. “∼” was used because the two halves have similar (but not exactly the same) sample sizes that summed to 5260 (total number of participants). Note that mass univariate associations (tstats) were computed independent of regression models and are therefore the same across Figures 1, 2, S1 and S2. Overall, across different sample sizes and behavioral domains, Haufe-transformed weights were more reliable than mass univariate associations (tstats), which were in turn more reliable than original regression weights.

### 3.2 Haufe-transformed weights are highly consistent across prediction models

The previous section investigated the reliability of feature importance across different data samples. Here, we seek to examine the reliability of feature importance across different prediction models in the full sample of 5260 participants. For each split-half of the 5260 participants, we computed the similarity (Pearson’s correlation) of feature importance across the prediction models.

Figure 3 shows the similarity of feature importance across prediction models. Consistent with Tian and Zalesky (2021), we found that Haufe-transformed weights showed better consistency than the original regression weights. Unlike Tian and Zalesky (2021), because of our significantly larger sample size, excellent consistency was observed for the Haufe-transformed weights (max = 0.97, min = 0.63). Interestingly, although random forests have rather different inductive biases from linear models, the Haufe-transformed weights of the random forests still exhibited strong similarity with the linear models, especially when predicting cognition.

**Figure 3.**
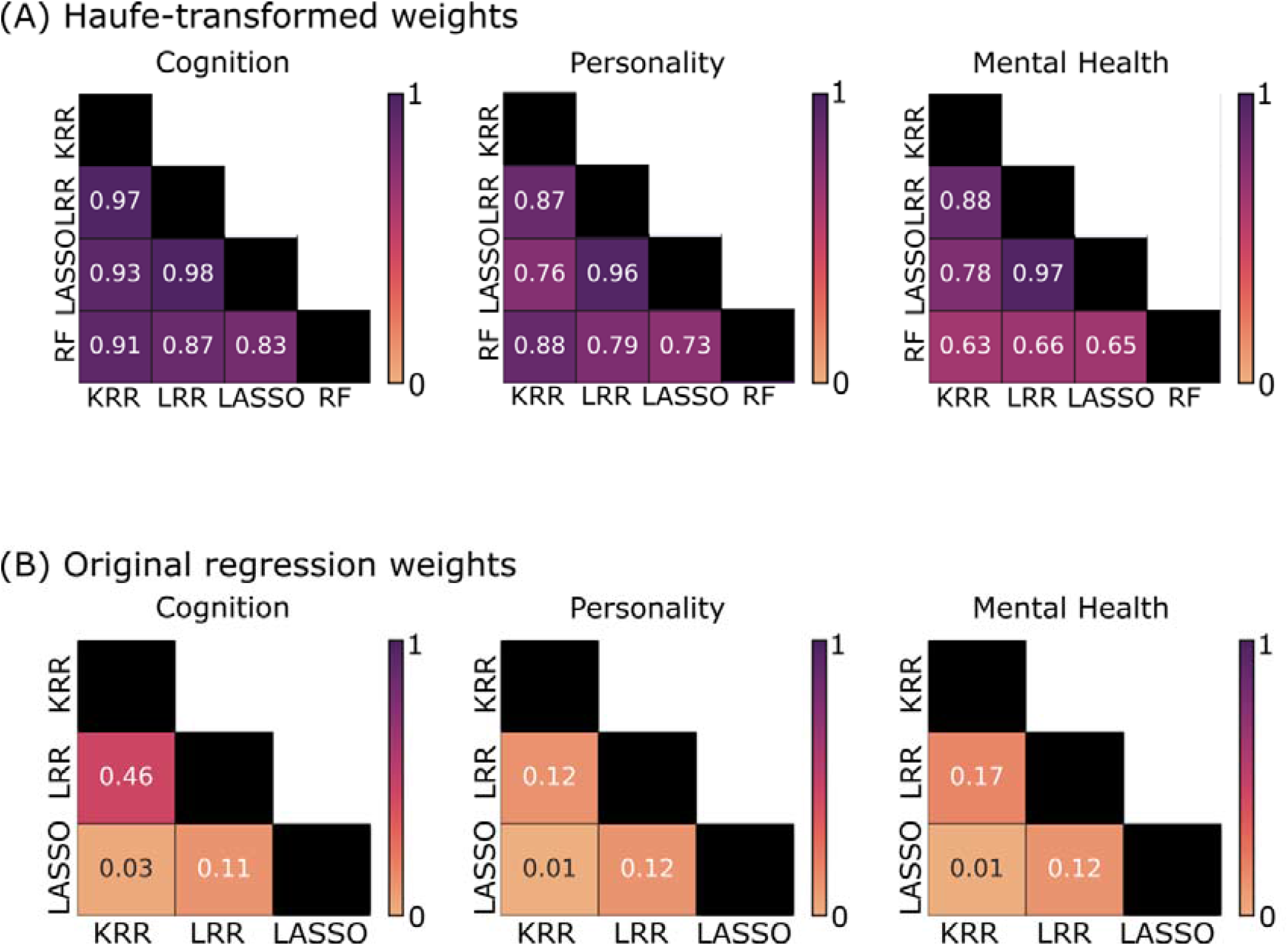
Similarity of feature importance across three predictive models in the full sample of 5260 participants. (A) Consistency of feature importance for Haufe-transformed weights. (B) Consistency of feature importance for original regression weights. We note that RF did not have regression weights, so did not appear in panel B. Similarity was computed as the Pearson’s correlation between feature importance values across different predictive models (KRR, LRR, LASSO and RF). Similarity was computed for each split-half and then averaged across the 126 data splits. Excellent consistency was observed for the Haufe-transformed weights.

### 3.3 Feature importance reliability is strongly positively correlated with prediction accuracy across behavioral measures

So far, our results have been largely consistent with Tian and Zalesky (2021), except our larger sample sizes led to better split-half reliability of the Haufe-transformed weights. Next, we investigated the relationship between prediction accuracy and split-half reliability of feature importance using the full sample of 5260 participants.

Split-half reliability and prediction accuracy of each behavioral score were computed for each split-half of the dataset, followed by averaging across the 126 data splits. Figure 4A shows the correlation between feature importance reliability and prediction accuracy across the 36 behavioral measures for KRR. Prediction accuracy was highly correlated with split-half reliability of Haufe-transformed model weights (r = 0.78), t-statistics (r = 0.94) and original regression weights (r = 0.97). This suggests that a behavioral measure that was predicted with higher accuracy also enjoyed better feature importance reliability.

**Figure 4.**
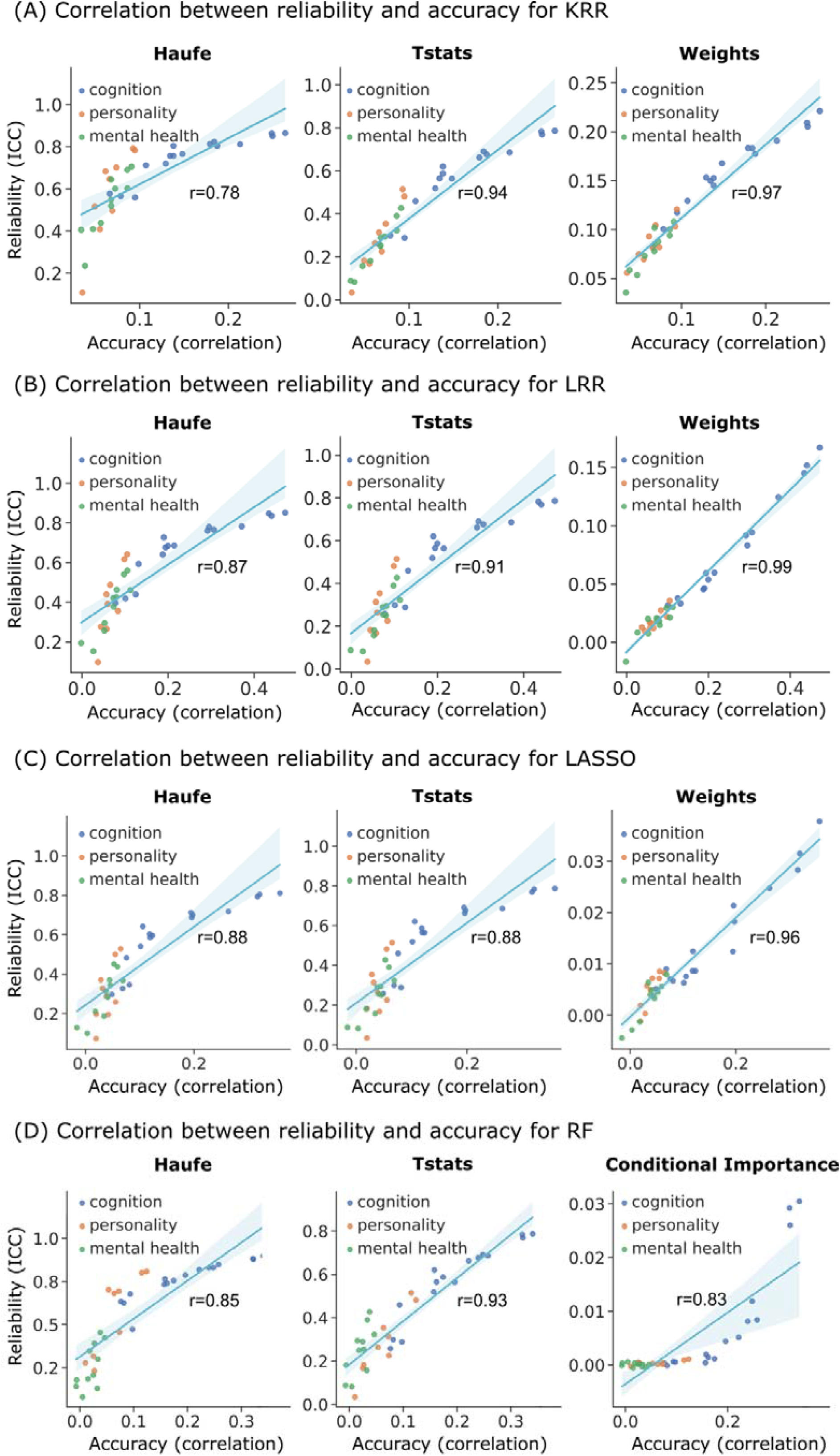
Split-half reliability of feature importance is positively correlated with prediction accuracy across 36 behavioral measures for (A) kernel ridge regression (KRR), (B) linear ridge regression (LRR), (C) LASSO and (D) random forest (RF). Split-half reliability and prediction accuracy of each behavioral score were computed for each split-half of the dataset, followed by averaging across the 126 data splits.

Similar conclusions were obtained with linear ridge regression (Figure 4B), LASSO (Figure 4C) and RF (Figure 4D). Overall, we found a strong positive relationship between feature importance reliability and prediction accuracy. We repeated the analysis with the behavioral component scores from Ooi et al (2022) and found similar positive relationships (Figure S3).

Furthermore, in the case of Haufe transform and univariate associations (t-stats), there appears to be a nonlinear relationship between prediction accuracies and split-half ICC (Figure 4). More specifically, higher accuracies led to greater split-half ICC, but with diminishing returns for behavioral measures with higher accuracies.

### 3.4 No clear relationship between prediction accuracy and feature importance reliability across predictive models

Table 1 summarizes average prediction accuracies for different behavioral domains, as well as split-half ICC of feature importance using the full sample of 5260 participants.

**Table 1.**
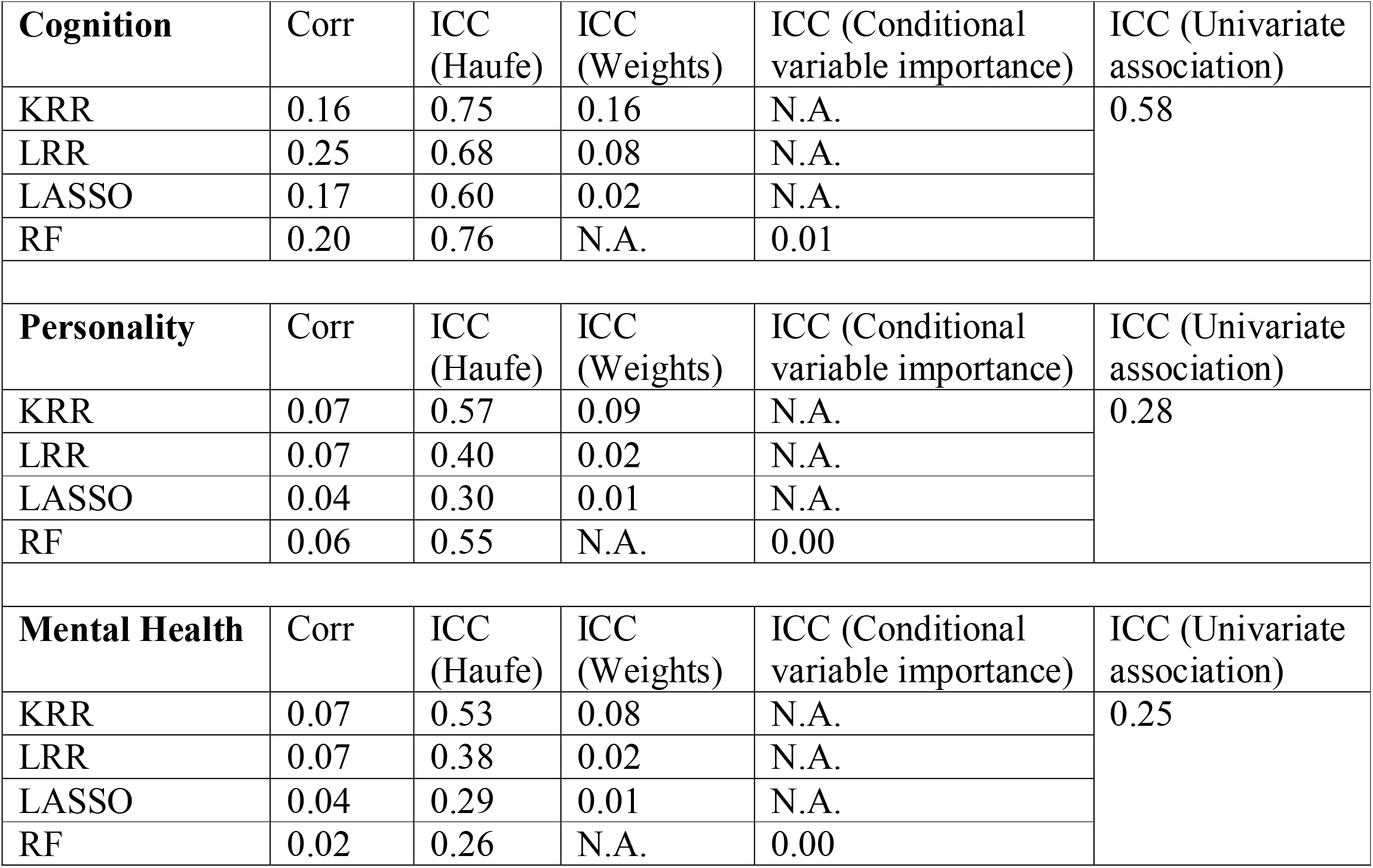
Summary of average prediction performance for cognitive, personality and mental health measures, as well as split-half ICC of Haufe-transformed weights, original weights, conditional variable importance and univariate associations (t-statistics). In general, within a behavioral domain (e.g., cognition), lower (or higher) prediction performance for a given predictive model was not necessarily associated with lower (or higher) split-half ICC.

Among the linear models, KRR exhibited the highest split-half ICC, but not necessarily the best prediction performance. LASSO generally had the worse prediction performance and the worst split-half ICC. LRR exhibited the best prediction performance, but an intermediate level of split-half ICC. On the other hand, the RF models exhibited good prediction performance and Haufe-transform split-half ICC for cognition and personality, but not for mental health. Overall, there was no clear relationship between prediction performance and feature importance reliability.

Note that in our other studies (Chen *et al*. 2022, Ooi *et al*. 2022), the prediction performance of KRR was similar to (or slightly better) than LRR, suggesting that depending on the dataset (or even across different samples within the same dataset), prediction accuracies can vary across prediction approaches.

### 3.5 Split-half reliability is necessary, but not sufficient, for correct feature importance

We have shown a strong positive correlation between feature importance reliability and prediction accuracy (Figure 4). There is also a lack of relationship between prediction accuracy across prediction models and feature importance reliability (Table 1). In the remaining sections of this study, we will delve more deeply into the mathematical relationships among feature importance reliability, feature importance error and prediction error.

We begin by showing that split-half feature importance reliability is necessary but not sufficient for obtaining the “correct” feature importance. Let *f*_*G*_ be the hypothetical ground-truth feature importance that might be derived assuming the correct generative process relating brain features and behavioral measures is known. However, in the following analysis, we do not assume the ground truth generative process is known and we make no assumption about how *f*_*G*_ can be computed even if the ground truth generative process is known.

Let *f*_*S*_ be the feature importance estimated from data sample *S*. Both *f*_*G*_ and *f*_*S*_ are *D* × 1, where *D* is the number of features. The expected feature importance error can be defined as the expectation of the squared error across different data samples *S*: *E*_*S*_ [(*f*_*G*_ − *f*_*S*_)^*T*^ (*f*_*G*_ − *f*_*S*_)]. Let 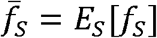 be the feature importance averaged across all possible data samples *S*. The feature importance error can then be decomposed into two terms:

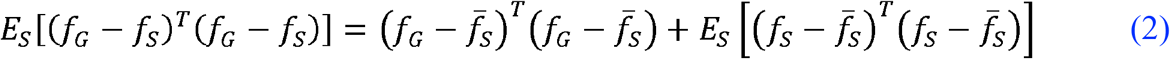

The proof is provided in the Appendix A. The decomposition of feature importance error as in Eq. (2) is similar in spirit (and derivation) to the classical bias-variance decomposition of prediction error.

The first term 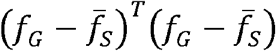 in Eq. (2) measures the bias of the feature importance estimation procedure. The second term 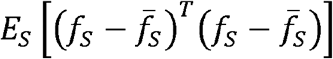 measures the variance of the estimated feature importance across different samples, which is the opposite of reliability. In other words, higher variance in feature importance estimation is the same as lower reliability. Therefore, from Eq. (2), we note that low feature importance variance (i.e., high feature importance reliability) is necessary but not sufficient for low feature importance error. Low feature importance variance must be coupled with low feature importance bias to achieve a small feature importance estimation error.

### 3.6 Prediction error reflects feature importance error for linear models

The previous section shows that the reliability of feature importance is not sufficient for low feature importance error. In this section, we show that when the ground truth data generation model is linear and feature importance is defined as regression weights (or Haufe-transformed weights), then the prediction error is directly related to the feature importance error.

A linear regression model assumes that the data is generated through a linear combination of features. For example, assume that a given data point (*x*_*i*_, *y*_*i*_) is generated by a linear model 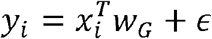. Here, *y*_*i*_ is a scalar, *x*_*i*_ is a *D* × 1 vector, *w*_*G*_ is the groundtruth *D* × 1 regression weights, and *D* is the number of features. *ϵ* is an independent noise term with zero mean. Without loss of generality, we assume that the expectation *y* of across data samples is 0 and the expectation of *x* across data samples is 0 for every feature. In the case of FC prediction of behavioral traits, each data sample is a participant.

Suppose data sample *S* = {(*x*_1_, *y*_1_), … (*x*_*N*_, *y*_*N*_)} is drawn as the training set. We can then train a linear regression model (e.g., LRR or LASSO) on *S* and obtain the regression weights *w*_*S*_. The resulting prediction model will be *ŷ* = *x*^*T*^*w*_*S*_. Let the difference between the ground truth and estimated weights be Δ_*w*_(*S*) = *w*_*G*_ − *w*_*S*_. Thus, the regression weights error (on average across different training sets *S*) can be defined as *E*_*S*_ [(*w*_*G*_ − *w*_*S*_)^*T*^ (*w*_*G*_ − *w*_*S*_)] = *E*_*S*_[Δ_*w*_(*S*)^*T*^ Δ_*w*_(*S*)].

On the other hand, the expected prediction error of the prediction algorithm can be defined as *E*_*S*_ *Ex,y* [(*y* − *x*^*T*^*w*_*S*_)^2^]. Here, *E*_*x,y*_ is the expectation of the squared prediction error over out-of-sample test data points sampled from the distribution of (*x, y*). We note that the test data points are sampled independently from the sampling of the training dataset. Then, the expected test error can be decomposed into:

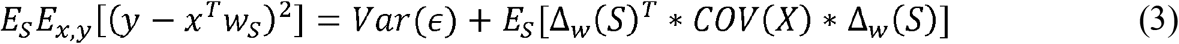

The proof is found in the Appendix B. In Eq. (3) the first term is the irreducible error *Var* (*ϵ*), which is the variance of the noise. The second term *E*_*S*_[Δ_*w*_(*S*)^*T*^ * *COV* (*X*) * Δ_*w*_(*S*)] is determined by both the regression weights error Δ_*w*_(*S*) and the covariance of features *COV* (*X*).

We can consider three different scenarios for the covariance matrix *COV* (*X*). First, suppose *COV* (*X*) is an identity matrix, which implies the features are independent and of unit variance. Then, the prediction error (Eq. (3)) can be written as *Var* (*ϵ*) + *E*_*S*_ [Δ_*w*_(*S*)^*T*^ Δ_*w*_(*S*)]. Therefore, the prediction error is simply the sum of the regression weights error and the irreducible error.

Second, suppose *COV* (*X*) is a diagonal matrix, i.e., *COV* (*X*) = *diag* (*σ*_1_, *σ*_2_, …, *σ*_*d*_), which implies the features are independent. In this case, the prediction error (Eq. (3)) can be written as 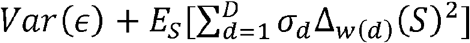. Here, Δ_*w*(*d*)_(*S*)is the regression weight error of the *d*-th feature based on the training dataset *S*. In this scenario, a bigger regression weights error still leads to a bigger prediction error, but the weights error of features with larger variance results in a larger prediction error than features with small variance.

Third, suppose we do not make any independence assumptions about the features. Since *COV* (*X*) is a symmetric matrix, we can decompose *COV* (*X*) as *COV* (*X*) = *R*^*T*^*DR*. Here, *R* is a rotation matrix where *R*^*T*^*R* is equal to an identity matrix and *D* is a diagonal matrix. Then, we can rewrite the prediction error (Eq. (3)) as:

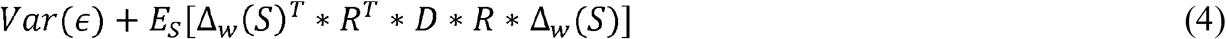

To summarize the three scenarios for *COV* (*X*), regression weights errors of all features are related to prediction error, but features with a larger variance (up to a rotation) have a stronger relationship with the prediction error.

We can also establish a similar relationship between the Haufe-transformed weights error and the prediction error. Note that the Haufe-transformed weights can be computed as *COV* (*X*_*S*_) * *w*_*s*_. Here the *w*_*s*_ is the original regression weights and *COV* (*X*_*S*_) is the feature covariance of training sample S. Assuming that the sample covariance is close to the true covariance, i.e., *COV* (*X*_*S*_) ≈ *COV* (*X*), then the Haufe-transformed weights error can be written as:

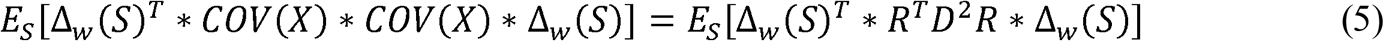

Comparing the Haufe-transformed weights error (Eq. (5)) with the prediction error (Eq. (4)), we see that the Haufe-transformed weights error is closely related to the prediction error, given that Eq. (4) and Eq. (5) only differ by the square of the diagonal matrix *D*.

Overall, we conclude that higher original regression weights errors (Eq. (4)) and higher Haufe-transformed errors (Eq. (5)) are related to greater prediction error up to a scaling by the feature covariance matrix.

## 4. Discussion

We have provided empirical and theoretical evidence on the relationship between prediction accuracy and feature importance reliability.

### 4.1 Haufe-transformed model weights are more reliable than original regression weights and univariate FC-behavior correlations

Consistent with Tian and Zalesky (2021), we found that Haufe-transformed weights were much more reliable than original regression weights. In our experiments, we note that even with a sample size of ∼2630 participants, the original kernel regression weights achieved a split-half ICC of less than 0.2 when predicting cognitive measures, which is less than the split-half ICC of Haufe-transformed weights with a sample size of 200 (Figure 1A). This is perhaps not surprising since it has been empirically shown that regression weights contain more noise than the Haufe-transformed weights (Haufe *et al*. 2014). Furthermore, for predictive models with sparse regularization (e.g., LASSO), it is well-known that noise in the features can lead to very different features being selected, which will lead to low split-half reliability in the regression weights. In the case of random forests, the poor split-half reliability was probably due to the large number of functional connectivity features (87571 features). Using a random forest with 100 trees and a depth of 4 would mean that a maximum of 1500 unique features being chosen, so the conditional variable importance was susceptible to the random choice of features included in the random forest.

Also consistent with Tian and Zalesky (2021), we found that Haufe-transformed weights were more reliable than univariate brain-behavior correlations. In our experiments, we note that with a sample size of ∼2630 participants, the univariate FC-behavior correlations achieved a split-half ICC of less than 0.6 for cognitive measures, which is less than the split-half ICC of Haufe-transformed weights with a sample size of 1000 (Figure 1A). The higher split-half ICC of Haufe-transformed weights over univariate associations is somewhat surprising. A previous study has suggested that the predicted outcomes of predictive models is substantially more reliable than the functional connectivity features themselves (Taxali *et al*. 2021). Here, we speculate that the predicted behavioral measures might even be more reliable than the raw behavioral measures themselves. The reason is that the regularization of many predictive models serves to “shrink” the predicted outcomes towards the population mean, which should increase reliability. If predicted behavioral measures are more reliable than raw behavioral measures, then the covariance of the predicted behavioral measures with FC (i.e., haufe-transformed weights) should be more reliable than the correlation between raw behavioral measures and FC (i.e., univariate associations).

It is also worth mentioning that Tian and Zalesky (2021) found that the split-half ICC of Haufe-transformed weights remained lower than 0.4 across split-half of 800 participants (i.e., two groups of 400 participants), which is consistent with our results (see sample size of 400 in our Figures 1 and 2). Not surprisingly, we obtained higher reliability with larger sample sizes. More specifically, with a sample size of about 2600 participants, Haufe-transformed weights achieve average intra-class correlation coefficients of 0.75, 0.57 and 0.53 for cognitive, personality and mental health measures respectively (Figure 1). Overall, the use of Haufe-transformed weights might help to alleviate reliability issues highlighted in previous neuroimaging studies (Kharabian Masouleh *et al*. 2019, Marek *et al*. 2022). On the other hand, we recommend that regression weights should not be used for model interpretation given their low split-half reliability even in the large sample regime of a few thousand participants.

### 4.2 There is not always an empirical trade-off between feature importance reliability and prediction accuracy

Tian and Zalesky (2021) found that FC-based prediction using lower resolution atlases (compared with higher resolution atlases) had higher feature importance reliability but lower prediction accuracy. Our study suggests that this trade-off between prediction accuracy and feature importance reliability is not universal. For example, we found that behavioral measures that are predicted better also enjoy better feature importance reliability (Figure 4).

Furthermore, in our current study, within a behavioral domain, there was no clear relationship between prediction performance and feature importance reliability across regression algorithms (Table 1). Similarly, as can be seen in Figure 2 of Tian and Zalesky (2021), higher prediction accuracy does not necessitate lower split-half reliabilities, e.g., kernel ridge regression enjoyed better prediction accuracy *and* feature importance reliability than connectome-based predictive modeling.

Overall, these empirical results show that it is possible to achieve high prediction accuracy *and* high feature importance reliability, suggesting that there is not always a trade-off between prediction accuracy and feature importance reliability.

### 4.3 There is not a theoretical trade-off between feature importance reliability and prediction accuracy

Eq. (2) shows that feature importance reliability is necessary but not sufficient for obtaining the “correct” feature importance (or low feature importance error). More specifically, feature importance error can be decomposed into a bias term and a variance term, where the variance term is the opposite of feature importance reliability. Consequently, low feature importance variance (i.e., high feature importance reliability) is necessary but not sufficient for low feature importance error.

This result echoes previous studies in neuroimaging (Noble *et al*. 2017), as well as other areas of quantitative research (Kirk and Miller 1986), demonstrating that reliability is not the same as validity. To give an extreme example, if we utilized an extremely strong regularization in our regression models, the regression weights would be driven to zero. In this scenario, the feature importance (regression weights) would be highly reliable across data samples, but the feature importance would not be valid or close to the ground truth values (derived from the ground truth generative process).

In the case of linear models, we further showed in Eq. (3) that higher feature importance error (operationalized by original regression weights) is related to worse prediction accuracy, up to a rotation and scaling by the feature covariance matrix. In Eq. (5), we showed that higher feature importance error (operationalized by Haufe-transformed weights) is related to worse prediction accuracy, up to a scaling of the eigenvalues of the feature covariance matrix.

Overall, these theoretical results suggest that at least in the case of linear models, there is not necessarily a trade-off between feature importance reliability and prediction accuracy. In fact, improving prediction performance might even reduce feature importance error and potentially improve feature importance reliability.

Given the link between feature importance error and prediction performance, in some sense, feature importance error might be a more meaningful metric than feature importance reliability. However, we cannot directly measure feature importance error, so a useful proxy might be to consider both feature importance reliability and prediction performance.

### 4.4 Reliability of functional connectivity and behavioral measures

There is a significant literature on the reliability of FC (Noble *et al*. 2019). Recent studies have also emphasized that the reliability of behavioral measures (in addition to FC reliability) is important for good prediction performance (Nikolaidis *et al*. 2022, Gell *et al*. 2023). How do FC and behavioral reliability relate to our theoretical results?

Recall that assuming *y* = *x*^*T*^*w*_*G*_ + *ϵ*, then as shown in Eq. (3), the prediction error can be written as *Var* (*ϵ*) + *E*_*S*_ [Δ_*w*_(*S*)^*T*^ * *COV* (*X*) * Δ_*w*_(*S*)], where *COV* (*X*) is the *D* × *D* feature covariance matrix and Δ_*w*_(*S*) is the *D* × 1 regression weights error. *Var* (*ϵ*) is the variance of the irreducible noise *ϵ*, which can be thought of as the variance of the behavioral measure unrelated to the features.

To think about the effect of the reliability of behavioral measure y, suppose we add more noise to the behavioral measure *y*, so that *y* = *x*^*T*^*w*_*G*_ + *ϵ*_2_, where *Var* (*ϵ*_2_) > *Var* (*ϵ*). In this scenario, the prediction error can now be written as *Var* (*ϵ*_2_) + *E*_*S*_ [Δ_*w*_(*S*)^*T*^ * *COV* (*X*) * Δ_*w*_(*S*)]. Therefore, the equation is basically the same as before except for the larger noise variance *Var* (*ϵ*_2_). In addition, the larger noise in *y* would also lead to greater regression weights error Δ_*w*_(*S*). Overall, worse behavioral reliability leads to larger *Var* (*ϵ*) and Δ_*w*_(*S*), and thus worse prediction error. The worse regression weights error Δ_*w*_(*S*) is in turn associated with worse feature importance bias and/or reliability (via Eq. (2)).

On the other hand, suppose we add more noise to the FC features *x* to reduce FC reliability, so that the new features *x*_2_ = *x* + *ϵ*_2_, where *ϵ*_2_ is a *D* × 1 noise vector with zero mean. Then 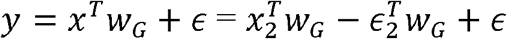. Using the same derivation as Appendix B, the prediction error can now be decomposed as 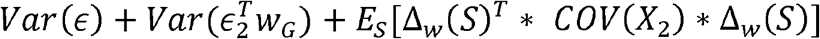. Therefore, worse FC reliability leads to an additional noise term 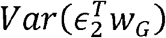, larger feature covariance matrix *COV* (*X*_2_) and greater regression weight error Δ_*w*_(*S*), thus leading to worse prediction performance. The worse regression weights error Δ_*w*_(*S*) is in turn associated with worse feature importance bias and/or reliability (via Eq. (2)).

It is important to note that perfect feature and behavioral reliability is not a panacea. For example, if features and behavioral measure both have perfect reliability, but they are not related to each other, e.g., *x* and *y* both follow white Gaussian distributions. Then, *y* = *x*^*T*^0 + *ϵ* (i.e., *w*_*G*_ = 0), and the prediction error of still follows Eq. (3): *Var* (*ϵ*) + *E*_*S*_ [Δ_*w*_(*S*)^*T*^ * *COV* (*X*) * Δ_*w*_(*S*)]. In this case, *Var* (*ϵ*)is simply the variance of the behavioral measure. We note that if we simply predict the mean of the behavioral measure (i.e., completely ignore *x*), then the prediction error of *y* will be *Var* (*ϵ*). On the other hand, if we are fitting a model to predict *y* from *x*, then because of the finite sample size, the regression weights will not be equal to the ground truth, so the regression weight error Δ_*w*_(*S*) is actually non-zero. Therefore, the overall prediction will be worse than simply predicting the mean of the behavioral measure, which would lead to a negative coefficient of determinant (a measure of prediction performance).

### 4.5 Reconciling theoretical and empirical results

Our theoretical results suggest a link between feature importance reliability and prediction performance.

Consistent with the theoretical results, there was empirically a strong correlation between feature importance reliability and prediction performance across behavioral measures (Figure 4). Similar to our previous studies (Kong *et al*. 2021, Chen *et al*. 2022, Ooi *et al*. 2022), cognitive measures were predicted better than other behavioral measures (Figures 1, 2 and 4). One possible explanation for the variation in prediction performance across behavioral measures might be the reliability of the behavioral measures, as discussed in Section 4.4 and previous studies (Nikolaidis *et al*. 2022, Gell *et al*. 2023). Another possible explanation is the strength of the relationship between FC features and target behavioral measures (again discussed in Section 4.4). Therefore, behavioral measures with higher reliability and/or stronger relationship with FC features might be predicted better, as well as enjoyed better feature importance error and reliability.

On the other hand, there was empirically not a clear relationship between prediction performance and feature importance reliability across predictive models (Table 1). For example, when predicting cognition, KRR exhibited worse prediction performance than LRR (0.16 versus 0.25), but better Haufe-transformed feature importance reliability (0.75 vs 0.68). There are several ways these empirical and theoretical results can be reconciled.

First, although KRR exhibited better Haufe-transformed feature importance reliability than LRR, it is possible that KRR had worse feature importance bias than LRR, so that the overall feature importance error is worse than LRR, resulting in worse prediction performance.

Second, recall that according to Eq.(5), the prediction error can be expressed as *E*_*S*_ [Δ_*w*_(*S*)^*T*^ * *COV* (*X*) * Δ_*w*_(*S*)], where Δ_*w*_(*S*) is the *D* × 1 feature importance error (where *D* is the number of features) and *COV* (*X*) is the *D* × *D* covariance matrix of the features. Because of the middle covariance term, not all feature importance errors are equally important. It is possible that KRR has lower feature importance errors on average across all features (Δ_*w*_(*S*)^*T*^ Δ_*w*_(*S*)) than LRR, but the feature importance error is greater for certain features that are more intrinsically linked to the prediction error via the covariance term *COV* (*X*) + *COV* (*X*).

Third, Eq. (5) assumes that the true data generation process is linear, i.e., there is a linear relationship between FC features and the target variable. Therefore, Eq. (5) might not hold if the true relationship between FC features and target variable is nonlinear.

Finally, the prediction error in Eq. (5) is the average across infinite instances of training set S and an infinite test set. Therefore, the equation can be violated in the finite sample scenario (Table 1).

## 5. Conclusion

Firstly, we show that Haufe-transformed weights are much more reliable than original regression weights when computing feature importance. Secondly, feature importance reliability is strongly positively correlated with prediction accuracy across phenotypes. However, within a particular behavioral domain, there is no clear relationship between prediction performance and feature importance reliability across regression models. Thirdly, we show mathematically that feature importance reliability is necessary, but not sufficient, for low feature importance error. In the case of linear models, lower feature importance error is mathematically related to lower prediction error. Therefore, higher feature importance reliability might yield lower feature importance error and higher prediction accuracy. Finally, we discuss how our theoretical results relate with the reliability of imaging features and behavioral measures.

## Acknowledgements

Our research is currently supported by the Singapore National Research Foundation (NRF) Fellowship (Class of 2017), the NUS Yong Loo Lin School of Medicine (NUHSRO/2020/124/TMR/LOA), the Singapore National Medical Research Council (NMRC) LCG (OFLCG19May-0035), NMRC STaR (STaR20nov-0003), and the USA NIH (R01MH120080). Our computational work was partially performed on resources of the National Supercomputing Centre, Singapore (https://www.nscc.sg). Any opinions, findings and conclusions or recommendations expressed in this material are those of the authors and do not reflect the views of the Singapore NRF or the Singapore NMRC.

Data used in the preparation of this article were obtained from the Adolescent Brain Cognitive Development^SM^ (ABCD) Study (https://abcdstudy.org), held in the NIMH Data Archive (NDA). This is a multisite, longitudinal study designed to recruit more than 10,000 children age 9-10 and follow them over 10 years into early adulthood. The ABCD Study® is supported by the National Institutes of Health and additional federal partners under award numbers U01DA041048, U01DA050989, U01DA051016, U01DA041022, U01DA051018, U01DA051037, U01DA050987, U01DA041174, U01DA041106, U01DA041117, U01DA041028, U01DA041134, U01DA050988, U01DA051039, U01DA041156, U01DA041025, U01DA041120, U01DA051038, U01DA041148, U01DA041093, U01DA041089, U24DA041123, U24DA041147. A full list of supporters is available at https://abcdstudy.org/federal-partners.html. A listing of participating sites and a complete listing of the study investigators can be found at https://abcdstudy.org/consortium_members/. ABCD consortium investigators designed and implemented the study and/or provided data but did not necessarily participate in the analysis or writing of this report. This manuscript reflects the views of the authors and may not reflect the opinions or views of the NIH or ABCD consortium investigators. The ABCD data repository grows and changes over time. The ABCD data used in this report came from http://dx.doi.org/10.15154/1504041.

## Appendix A

In this appendix, we will provide proof of Eq. (2), which decomposes the feature importance error *E*_*S*_ [(*f*_*G*_ − *f*_*S*_)^*T*^ (*f*_*G*_ −*f*_*S*_)] into a bias term 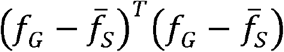 and a variance term 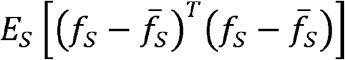.

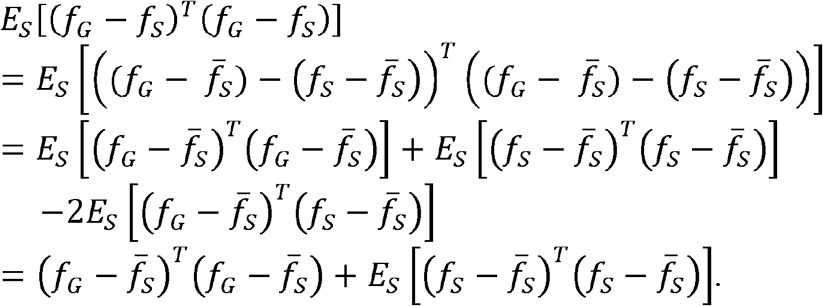

where the last equality is true because 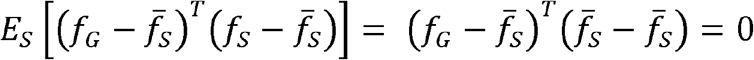.

## Appendix B

In this appendix, we will provide proof of Eq. (3), which establishes the relationship between the prediction error *E*_*S*_ *E*_*x,y*_[(*y* − *x*^*T*^*w*_*S*_)^2^] and regression weights error Δ_*w*_(*S*), assuming an underlying linear model 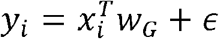:

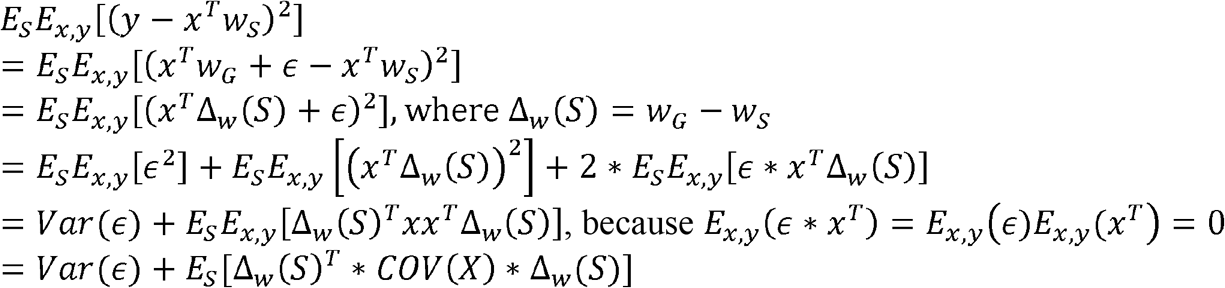

## Supplementary Materials

**Table S1.** Distribution of the included samples (n=5260) by site and scanner

| ABCD Site | Make | Model | N | Site-cluster |
| --- | --- | --- | --- | --- |
| 16 | Siemens | Prisma | 631 | A |
| 13 | GE | Discovery MR750 | 389 | B |
| 4 | GE | Discovery MR750 | 452 | C |
| 22 | GE | Discovery MR750 | 26 | C |
| 14 | Siemens | Prisma/Prisma fit | 287 | D |
| 15 | Siemens | Prisma fit | 190 | D |
| 6 | Siemens | Prisma fit | 311 | E |
| 9 | Siemens | Prisma fit | 201 | E |
| 10 | GE | Discovery MR750 | 403 | F |
| 11 | Siemens | Prisma | 228 | F |
| 3 | Siemens | Prisma | 356 | G |
| 5 | Siemens | Prisma fit | 183 | G |
| 2 | Siemens | Prisma fit | 220 | H |
| 7 | Siemens | Prisma fit | 172 | H |
| 8 | GE | Discovery MR750 | 184 | I |
| 20 | Siemens | Prisma/Prisma fit | 286 | I |
| 12 | Siemens | Prisma fit | 283 | J |
| 18 | GE | Discovery MR750 | 215 | J |
| 21 | Siemens | Prisma fit/Prisma | 251 | J |

**Figure S1.**
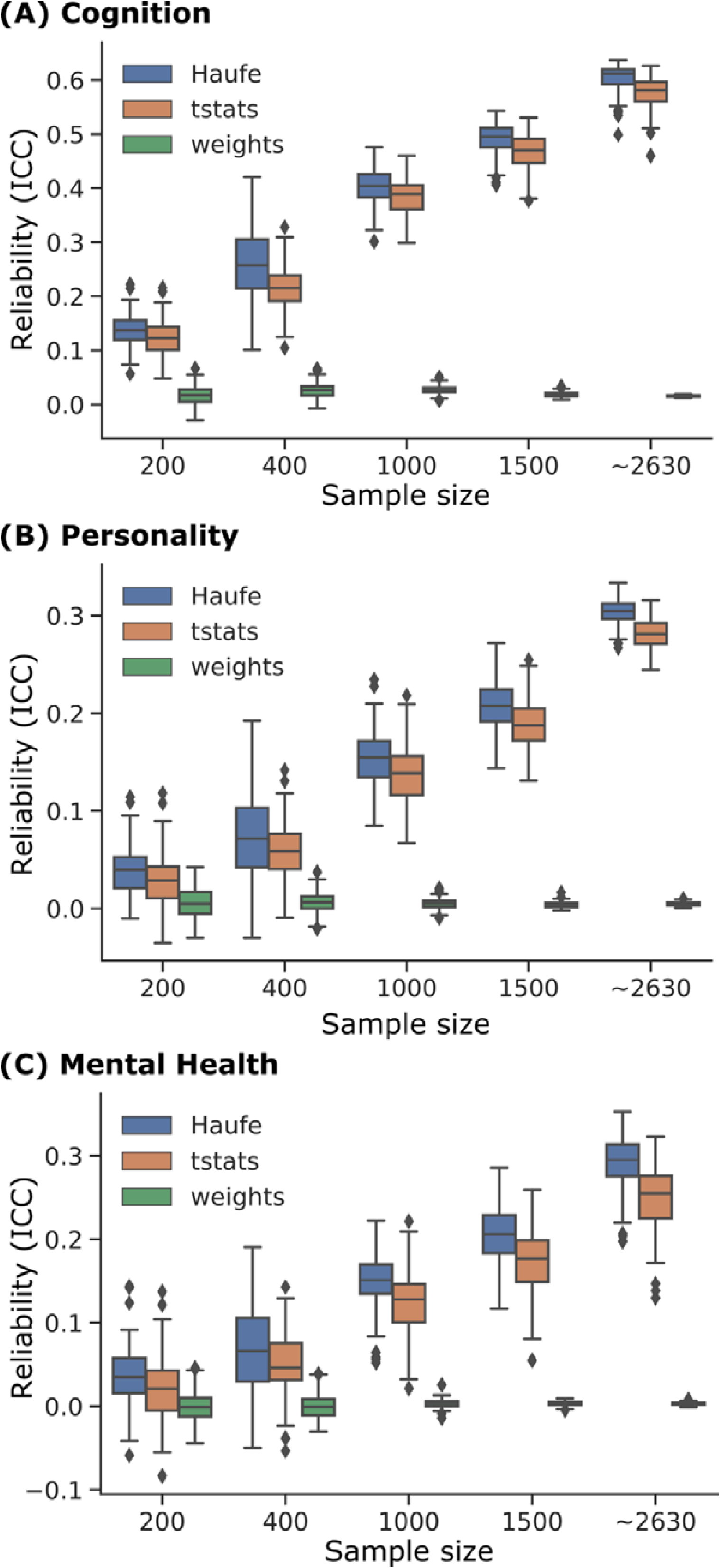
Split-half reliability of feature importance of LASSO models across different sample sizes, interpretation methods, and behavioral domains: (A) cognition, (B) personality, and (C) mental health. Same as Figure 1, except using lasso as the prediction model. Split-half reliability was computed as interclass correlation coefficients (ICC) of feature importance obtained from two non-overlapping split-halves of the dataset. After splitting, data were randomly subsampled to show the effect of sample size on feature importance reliability. Full data without subsampling was reported as a sample size of ∼2630. “∼” was used because the two halves have similar (but not exactly the same) sample sizes that summed to 5260 (total number of subjects). Note that BWA t-statistics (tstats) were computed independent of regression models and are therefore the same across Figures 1, 2, S1 and S2. Overall, across different sample sizes and behavioral domains, Haufe-transformed weights were more reliable than BWA t-statistics, which were in turn more reliable than regression weights.

**Figure S2.**
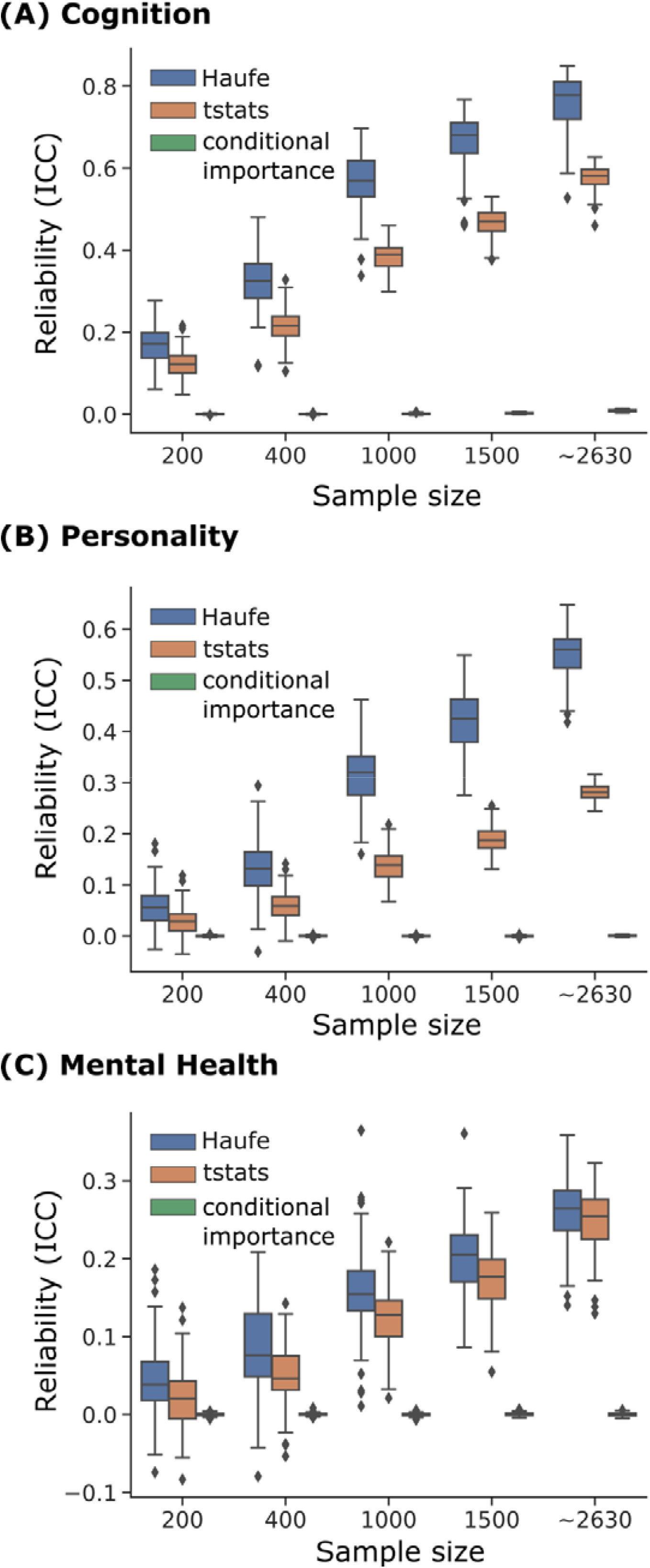
Split-half reliability of feature importance of RF models across different sample sizes, interpretation methods, and behavioral domains: (A) cognition, (B) personality, and (C) mental health. Same as Figure 1, except using random forest as the prediction model. Split-half reliability was computed as interclass correlation coefficients (ICC) of feature importance obtained from two non-overlapping split-halves of the dataset. After splitting, data were randomly subsampled to show the effect of sample size on feature importance reliability. Full data without subsampling was reported as a sample size of ∼2630. “∼” was used because the two halves have similar (but not exactly the same) sample sizes that summed to 5260 (total number of subjects). Note that BWA t-statistics (tstats) were computed independent of regression models and are therefore the same across Figures 1, 2, S1 and S2. Overall, across different sample sizes and behavioral domains, Haufe-transformed weights were more reliable than BWA t-statistics, which were in turn more reliable than conditional variable importance.

**Figure S3.**
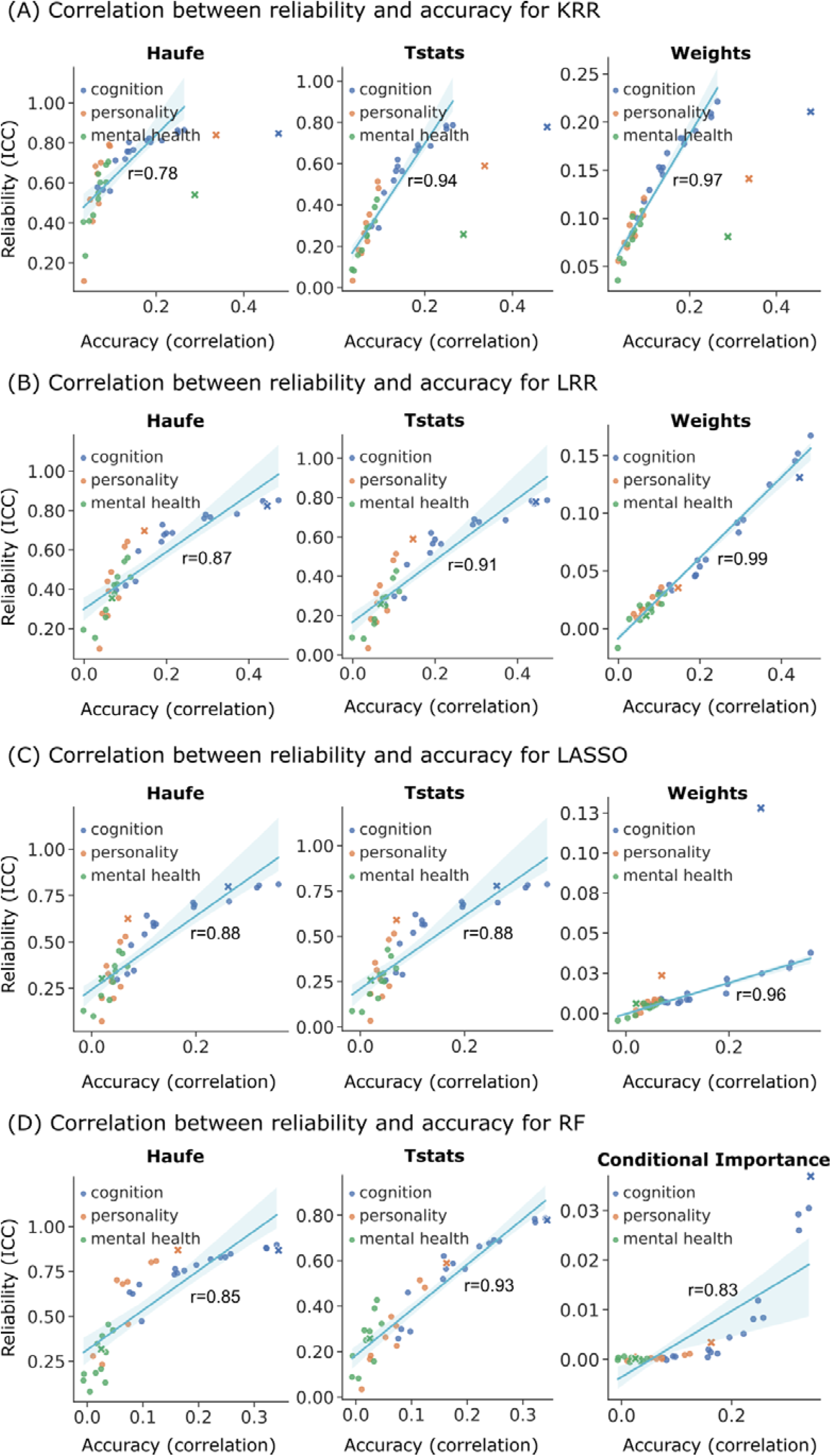
Split-half reliability of feature importance is positively correlated with prediction accuracy across 36 original behavioral measures (marked with circles) and 3 behavioral component scores (marked with crosses) for (A) kernel ridge regression (KRR), (B) linear ridge regression (LRR), (C) LASSO and (D) random forest (RF). Split-half reliability and prediction accuracy of each behavioral score were computed for each split-half of the dataset, followed by averaging across the 126 data splits. The results were highly similar to Figure 4, although in the case of KRR, the component scores were shifted to the right relative to the original regression line.

## Notes

### Competing Interest Statement

The authors have declared no competing interest.

### Summary of Updates

The manuscript has been revised to include a more detailed explanation of the Haufe transformation, additional analysis, and additional explanation of the theoretical results.

